# Universal Digital High Resolution Melt for the detection of pulmonary mold infections

**DOI:** 10.1101/2023.11.09.566457

**Authors:** Tyler Goshia, April Aralar, Nathan Wiederhold, Jeffrey D. Jenks, Sanjay R. Mehta, Mridu Sinha, Aprajita Karmakar, Ankit Sharma, Rachit Shrivastava, Haoxiang Sun, P. Lewis White, Martin Hoenigl, Stephanie I. Fraley

**Affiliations:** Department of Bioengineering, University of California San Diego, San Diego, CA, USA; Department of Pathology, University of Texas Health Science Center, San Antonio, TX, USA; Department of Medicine, Duke University School of Medicine, Durham, NC, USA; Durham County Department of Public Health, Durham, NC, USA; Department of Medicine, University of California San Diego, San Diego, CA, USA; San Diego Veterans Administration Medical Center, San Diego, CA, USA; MelioLabs, Inc., Santa Clara, CA, USA; Public Health Wales Microbiology Cardiff, and Cardiff University Centre for Trials Research/Division of Infection/Immunity, University Hospital of Wales, Cardiff, United Kingdom; Department of Medicine, Medical University of Graz, Graz, Austria

**Keywords:** IMI, dPCR, HRM, machine learning

## Abstract

**Background:** Invasive mold infections (IMIs) such as aspergillosis, mucormycosis, fusariosis, and lomentosporiosis are associated with high morbidity and mortality, particularly in immunocompromised patients, with mortality rates as high as 40% to 80%. Outcomes could be substantially improved with early initiation of appropriate antifungal therapy, yet early diagnosis remains difficult to establish and often requires multidisciplinary teams evaluating clinical and radiological findings plus supportive mycological findings. Universal digital high resolution melting analysis (U-dHRM) may enable rapid and robust diagnosis of IMI. This technology aims to accomplish timely pathogen detection at the single genome level by conducting broad-based amplification of microbial barcoding genes in a digital polymerase chain reaction (dPCR) format, followed by high-resolution melting of the DNA amplicons in each digital reaction to generate organism-specific melt curve signatures that are identified by machine learning.

**Methods:** A universal fungal assay was developed for U-dHRM and used to generate a database of melt curve signatures for 19 clinically relevant fungal pathogens. A machine learning algorithm (ML) was trained to automatically classify these 19 fungal melt curves and detect novel melt curves. Performance was assessed on 73 clinical bronchoalveolar lavage (BAL) samples from patients suspected of IMI. Novel curves were identified by micropipetting U-dHRM reactions and Sanger sequencing amplicons.

**Results:** U-dHRM achieved an average of 97% fungal organism identification accuracy and a turn-around-time of 4hrs. Pathogenic molds (*Aspergillus, Mucorales, Lomentospora* and *Fusarium)* were detected by U-dHRM in 73% of BALF samples suspected of IMI. Mixtures of pathogenic molds were detected in 19%. U-dHRM demonstrated good sensitivity for IMI, as defined by current diagnostic criteria, when clinical findings were also considered.

**Conclusions:** U-dHRM showed promising performance as a separate or combination diagnostic approach to standard mycological tests. The speed of U-dHRM and its ability to simultaneously identify and quantify clinically relevant mold pathogens in polymicrobial samples as well as detect emerging opportunistic pathogens may provide information that could aid in treatment decisions and improve patient outcomes.

## INTRODUCTION

Invasive mold infections (IMI) cause millions of infections globally and account for an estimated 1.6 million deaths annually(Bongomin et al. 2017). Patients at risk from IMIs, including both severely immunocompromised and also more immunocompetent individuals(Jenks et al. 2023), are increasing. IMIs in more immunocompetent persons/those receiving systemic corticosteroids are characterized by early tissue invasive growth in the lungs with bloodstream invasion potentially occurring later although not universally, while early angioinvasive growth is more common in severely immunocompromised persons (Jenks and Hoenigl 2018). Histopathologic examination and culture of tissue or bronchoalveolar lavage fluid (BALF) is considered the reference standard for IMI diagnosis, but is slow, with histopathology often available only at autopsy, while culture has poor sensitivity (Antinori, Corbellino, and Parravicini 2018). Incubation of fungal cultures for four weeks is considered best practice to maximize the recovery of slow growing species, with most detected by day 14 (Zhu and Qi, n.d.). BAL antigen tests, such as galactomannan (GM), can be helpful but are only positive for a limited number of specific mold organisms, and are further limited by variable turnaround times and lower sensitivity for individuals on mold-active antifungal prophylaxis or treatment (Jenks et al. 2023). The absence of rapid and accessible fungal diagnostics often results in empiric utilization of systemic antifungals, mostly targeted against *Aspergillus* spp., some of which are lacking activity against other molds (Denning et al. 2017). As a result, IMIs are often diagnosed and treated too late, leading to high mortality rates of 40-80%. It is estimated that 80% of patients could be saved with rapid diagnostics to inform early and targeted treatment (“Invasive Fungal Infections: A Creeping Public Health Threat” 2018).

Universal digital High-Resolution Melt (U-dHRM) to detect mold pathogens in BALF may be a promising hypothesis free, unbiased diagnostic approach that could achieve rapid and near point-of-care diagnosis to inform treatment decisions and improve patient outcomes. This approach consists of a single closed-tube test that integrates universal amplification of pathogen barcoding sequences in a digital polymerase chain reaction (dPCR) format with high resolution melting of DNA and machine learning (Fig. 1) (Langouche et al. 2021; Aralar et al. 2020; Sinha, Mack, et al. 2018; Velez et al. 2017). The integration and advancement of these techniques promises a unique combination of advantages: speed and breadth of detection, sensitivity and absolute quantification, and pathogen identification in polymicrobial samples(Sinha, Jupe, et al. 2018).

**Figure 1.**
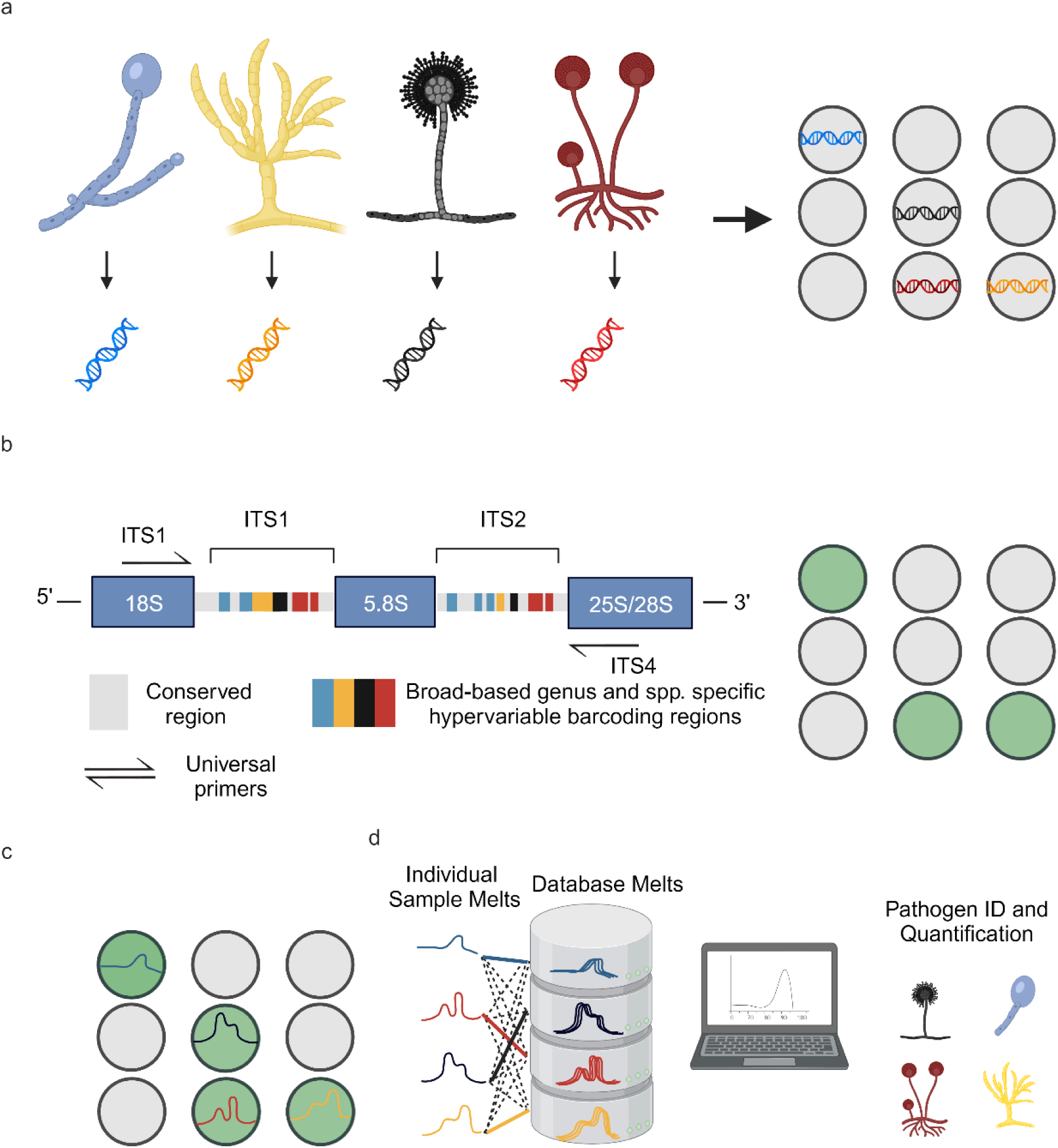
U-dHRM technology overview. a) Extraction of genomic DNA and digital loading. b) Universal amplification of fungal ITS barcoding region leading to a fluorescence increase in each positive reaction well. c) Barcode sequence-defined melt curve signatures. d) Automatic identification of each known pathogen melt curve and detection of novel melt curves using machine learning. Created with BioRender.com

Here, we advanced the U-dHRM assay and database for detection of IMI pathogens, advanced the machine learning algorithm to recognize database organism curves and also flag novel organism melt curves, and developed a dPCR reaction recovery method to Sanger sequence novel melt curves and expand the pathogen panel. We applied these advancements to test 75 clinical BALF samples, assessing the utility of this approach for IMI diagnosis compared to gold standard tests.

## METHODS

### ITS-Asp PCR

ITS1 (5’-TCCGTAGGTGAACCTGCGG -3’) and ITS4 (5’-TCCTCCGCTTATTGATATGC -3’) universal primers multiplexed with Asp 1 (5’-CGGCCCTTAAATAGCCCGGTC -3’) and Asp 2 (5’-ACCCCCCTGAGCCAGTCCG -3’) were used to amplify the Internal Transcribed Spacer (ITS) universal region for all fungi, and an Aspergillus specific region of 18S gene. ITS-Asp PCR was amplified in triplicates using the following protocol: each 15 µl reaction mixture contained 0.1 µM of each primer (IDT, Coralville, IA), 0.2 mM deoxynucleoside triphosphate (dNTP) (Invitrogen, Carlsbad, CA), 1X Phusion GC PCR buffer (Thermo Scientific, Waltham, MA), 2.5X EvaGreen (Biotium, Fremont, CA), 0.02 U/µl Phusion polymerase (New England Biolabs, Ipswich, MA), ultrapure water (Quality Biological, Gaithersburg, MD), and 3 µl of genomic DNA. Thermocycling for qPCR/dPCR and subsequent melt analysis were performed on a Quantstudio 3D real time PCR system and a ProFlex 2 x Flat Block Thermal Cycler (Applied Biosystems, Waltham, MA) using the QuantStudio 3D Digital PCR Chip (Applied Biosystems, Foster City, CA). The cycling conditions were as follows: hold at 98°C for 30 s, followed by 75 cycles of 98°C for 10 s, 61°C for 30 s, and 72°C for 60 s. At the end of cycling, there was a final extension step at 72°C for 5 min. PCR amplification was followed by a melt cycle of an initial denaturation at 95°C for 15 s, then heating from 65°C to 95°C.

### Control human β-actin PCR

Human beta actin primers F (5’-CGGCCTTGGAGTGTGTATTAAGTA -3’) and R (5’-TGCAAAGAACACGGCTAAGTGT -3’) were used to amplify the human β-actin gene. The PCR was amplified in triplicates using the following protocol: each 15 µl reaction mixture contained 0.1 µM each primers (IDT, Coralville, IA), 0.2 mM deoxynucleoside triphosphate (dNTP) (Invitrogen, Carlsbad, CA), 1X Phusion GC PCR buffer (Thermo Scientific, Waltham, MA), 2.5X EvaGreen (Biotium, Fremont, CA), 0.02 U/µl Phusion polymerase (New England Biolabs, Ipswich, MA), ultrapure water (Quality Biological, Gaithersburg, MD), and 3 µl of direct BALF sample liquid. Thermocycling for qPCR and subsequent melt analysis were performed on a Quantstudio 3D real time PCR system (Applied Biosystems, Waltham, MA). The cycling conditions were as follows: hold at 98°C for 30 s, followed by 55 cycles of 98°C for 10 s, 66°C for 30 s, and 72°C for 45 s. At the end of cycling, there was a final extension step at 72° for 5 min. PCR amplification was followed by a melt cycle of an initial denaturation at 95°C for 15 s, then heating from 65°C fo 95°C.

### Target DNA isolation

#### Melt curve database generation

The following fungal strains were provided as clinical isolates by Dr. Nathan Weiderhold at the Department of Pathology University of Texas Health Science Center, San Antonio, TX: *Aspergillus terreus, Aspergillus nidulans, Aspergillus versicolor, Mucor circinelloides, Mucor velutinosus, Mucor plumbeus, Rhizopus arrhizus var. delemar, Rhizopus microsporus, Lomentospora prolificans, Scedosporium apiospermum, Scopulariopsis brevicalus, Scopulariopsis candida,* and *Scopulariopsis gosspii. Aspergillus fumigatus, Aspergillus flavus, Aspergillus niger, Fusarium oxysporum, Cryptococcus neoformans, Candida krusei, Candida glabrata, and Candida albicans* were provided as clinical isolates from Dr. Sanjay Mehta at the San Diego VA Clinical Microbiology Laboratory. *Candida auris* was provided as a clinical isolate by Dr. Sharon Reed at the UCSD Center for Advanced Laboratory Medicine (CALM). For database generation, DNA was extracted using Lucigen MasterPure Yeast DNA Purification Kit (Lucigen, Middleton, WI, USA). DNA concentration was measured by bio-spectrophotometer absorbance readings and diluted to desired target concentrations.

#### Clinical samples host depletion and PCR inhibitor removal

Prior to DNA isolation, direct PCR β-actin was run to assess lavage quality as described previously. Clinical BALF sample DNA was isolated using Molysis Complete 5 Small Size Sample DNA Isolation (≤1ml Liquid) protocol (Molzym, Bremen, Germany). Each BALF sample was run in U-dHRM with Asp-ITS PCR conditions as described previously.

### Control pig BALF and analytical validation

For control and analytical spike-in experiments pig BALF was collected from euthanized pigs previously treated with antibiotics and anesthetized with ketamine/xylene/atropine. Ambu® aScope™ 4 Broncho single-use bronchoscopes (Ambu® A/S, Ballerup, Denmark) were used with 50 mL sterile isotonic irrigation 0.9% saline (NDC 0990-6138-22) for lavages. Prior to analytical validation, pig BALF was screened to be negative for target organisms by U-dHRM. Target organism spores were counted and plated, and six 10-fold serial dilutions were conducted to achieve concentrations down to 1 CFU/mL, with concurrent no spike controls. *A. fumigatus* and *C. albicans* spores from each concentration were spiked into 2 mL of pig BALF to achieve the final concentrations of 10k, 1k, 100, 10, 1 and 1 spores (CFU)/mL of BAL.

### DNA sequencing

The PCR products were prepared using ExoSAP-IT (Applied Biosystems, Foster City, CA) with the supplied protocol and then sent for Sanger sequencing (GENEWIZ, San Diego, USA) using the same respective forward primers in the 18S and ITS amplifications.

### Image processing and data analysis

A sequence of raw fluorescence images was captured during the heating and melting procedure for a single chip. Subsequently, these images underwent a sequence of image processing steps to identify and extract the individual wells within them along with their corresponding average intensity values. This procedure resulted in the translation of the average intensity measurements for each well across the entire set of images into a chronological array of values, thus creating a time series representation.

The original fluorescence time series, recognized as melt curves, underwent a two-fold transformation: initially, they were converted into their respective derivatives, after which they were subjected to a smoothing process using a Savitzky-Golay filter. Furthermore, these smoothed derivative time series were classified as ‘Positive’ if they exhibited a peak or local maxima beyond a temperature threshold of 85°C and with a minimum height of 4 units. In this context, a ‘Positive’ melt curve designates an instance where the presence of a particular fungal target is anticipated, whereas the remaining instances are categorized as ‘Negatives’.

Leveraging these identified ‘Positive’ melt curves, a dataset for machine learning purposes was constructed. Each time series within this dataset represented a derivative melt curve that was smoothed using a Savitzky-Golay filter with specific parameters: a window length of 9 and a polynomial order of 3. These time series were then normalized using Area Under the Curve Normalization to take care of scaling differences.

### Machine learning

We constructed a model based on our established database of organisms. This approach unfolds through a two-step procedure. The dataset we employed comprises a comprehensive set of 12,000 melt curves attributed to each distinct organism. Within this dataset, a subset of 10%, equating to 1,200 random melt curves, were selected and subjected to a time series DTW distance based K-means clustering process, yielding a culmination of up to 50 representatives (Lloyd n.d.) (MacQueen 1966) (“A Global Averaging Method for Dynamic Time Warping, with Applications to Clustering” 2011). Clusters housing fewer than 10 melt curves were excluded from consideration due to their susceptibility to noise-related interference. Owing to the substantial variability and inherent noise within the melt curves, we employed the K-means clustering technique as the initial step to disentangle pivotal clusters of variation, thereby yielding corresponding cluster centers that serve as robust and condensed representations of signals. These cluster centers are referred to as “DB representatives” in the flow charts in Fig. 3.

Another point to note is that instead of using the usual Euclidean distance-based K-means, we use Dynamic Time Warping (DTW) for both the cluster assignment as well as the averaging step of K-means. Temporal distortions (or shift) along the temperature (or time) axis causing well-to-well as well as chip-to-chip variations in melt curves is something inherent in HRM (Gundry et al. 2008) (Sinha, Mack, et al. 2018) (Langouche et al. 2021) and can be dealt with by using the various elastic distance measures for time series – amongst which the most popular one is Dynamic Time Warping (DTW) (Langouche et al. 2021; Sakoe and Chiba n.d.) and its variations (Rabiner, Rabiner, and Juang 1993) (“Weighted kNN and Constrained Elastic Distances for Time-Series Classification” 2020). More specifically, as we use DTW distance, we employ a more suitable DTW-based Barycenter Averaging (DBA) technique, as proposed by François et. al (“A Global Averaging Method for Dynamic Time Warping, with Applications to Clustering” 2011), for the K-means averaging step.

Subsequently, the second phase (Fig. 3) entailed the development of a classifier grounded in a 3 nearest neighbor (3NN) framework, leveraging the Euclidean distance as the defining metric. In this step, each test curve underwent alignment with every representative curve curated from the database (see blue boxes Fig. 3 a-b). Consequently, the KNN model was executed to discern the three nearest neighbors for each aligned test curve (see pink boxes Fig. 3 a-b) (Fix and Hodges 1951). The alignment procedure was deemed necessary to account for the potential shift-based discrepancies present among melt curves.

The outcome of this model furnishes predictions wherein concordance among the majority of neighbors designates a high-confidence classification. Conversely, instances in which all three nearest neighbors correspond to dissimilar organisms are categorized as low-confidence and consequently disregarded. Low-confidence instances can originate from either noisy signals or from novel curves that remain un-represented within the existing database. The performance of classification was quantified through the assessment of accuracy for each organism.

Although in literature the techniques novelty detection (ND), anomaly detection (AD) and outlier detection (OD) have been used interchangeably (“A Review of Novelty Detection” 2014). However, unlike AD or OD which usually refer to noisy or erroneous signals, ND usually has a positive learning attitude where the novel point is treated as a resource for potential future use (“A Review of Novelty Detection” 2014; Ruff et al. n.d.) (Pang et al. 2020). Currently AD, ND and OD are being studied under the common framework of Generalized Out of Distribution Detection (OOD) (Yang et al. 2021). Specifically for time series data, there is a significant amount of literature on AD but this research primarily focuses on finding point or subsequence anomalies within a large time series (Darban et al. 2022). As we have a larger number of smaller length time series we consider each time series (melt curve) as a separate data point. We then use a distance based OOD methodology for novelty detection (see section 5.3 of (Yang et al. 2021)) where the test curve is checked if it is outside of a certain standard deviations (threshold) away from each of the nearest three DB representatives (class cluster centers) obtained via 3NN step described previously. If this check is successful then the test point is certified as out of distribution and labeled as ‘novel’.

Furthermore, when dealing with patient samples, their time series were initially clustered utilizing the Euclidean-based K-means method (top left Fig. 3b). The resultant cluster centers were then subjected to classification leveraging the pre-constructed 3NN-based classifier designed for the database curves.

### Patients and Samples

In this retrospective case control study, banked BALF samples originated from patients with various underlying diseases and clinical suspicion of invasive pulmonary aspergillosis (IPA) or IMI, and galactomannan (GM) and *Aspergillus* spp. culture testing ordered between 2015 and 2019 at the University of California San Diego (UCSD). IMI was classified according to the revised European Organization for Research and Treatment of Cancer (EORTC)/Mycoses Study Group (MSG) criteria(Donnelly et al. 2019) and slightly modified AspICU criteria(Blot et al. 2012) (i.e., including positive BALF fluid GM of 1.0 ODI as entry criterion(Koehler et al. 2021)) (Meersseman et al. 2008) for patients in the intensive care unit (ICU) who did not fulfill EORTC/MSG host criteria. GM testing with the Platelia enzyme-linked immunosorbent assay (ELISA) (Bio-Rad Laboratories, Marnes-la-Coquette, France) was routinely and prospectively performed in all BALF samples before samples were stored at -70°C for up to 8 years. Based on classification we retrospectively tested 75 patient BALF samples from 30 patients classified as having suspected IPA with proven (n = 1), probable (n = 25), or putative (n = 4) IPA infections, and from 45 patients classified with limited evidence or as not having IPA (n=10 not classifiable, n = 4 possible IPA, n = 31 classified as no IPA). Not classifiable samples tested positive for mycological evidence and came from patients with clinical suspicion of IMI who did, however, not fulfill host factor criteria and/or did not present with typical radiological signs and were not admitted in the ICU. Direct β-actin PCR was used to access lavage quality according to previously published methods (Springer et al. 2018) (Lagier et al. 2016) (Fréalle et al. 2009). Two samples (n=1 possible and n=1 no IPA) were excluded due to no human DNA being detected.

### Novelty detection and Micromanipulator interrogation

Novel curves that were un-represented within the existing database were identified with ML as described previously. These curves’ physical X-Y on-chip coordinates were then identified using Melio Melt Inspector software (MelioLabs, Inc., Santa Clara, CA, USA). A custom micromanipulator setup then sampled the target amplicons from individual or clusters of wells using a glass capillary. Sampled amplicons were either reamplified with Asp-ITS primers or sent directly for Sanger sequencing. Reamplified Asp-ITS dPCR chips were used to demonstrate the process of adding novel organisms to the established database.

## RESULTS

### Fungal U-dHRM Assay Development and Analytical Validation

To develop a universal PCR assay for fungal detection, we first selected primers targeting conserved sequence regions flanking the ITS1-ITS4 barcoding region of the fungal genome (Supplementary Fig. 1) and tested their ability to amplify 21 clinically relevant organisms (Table 1). We started with *Aspergillus* spp., since it is the most prevalent IMI pathogen worldwide, and *Candida* spp., the most prevalent commensal genus, and began testing the ITS primers.

**Table 1.**
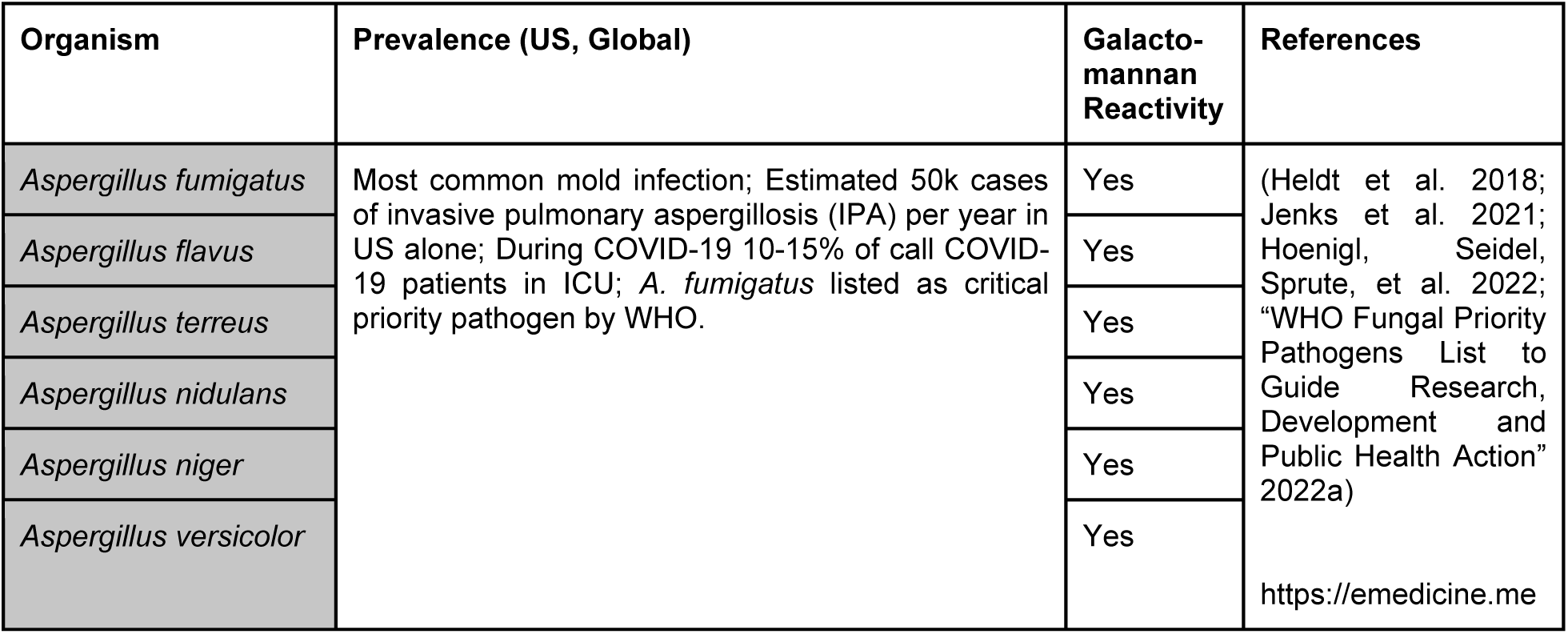
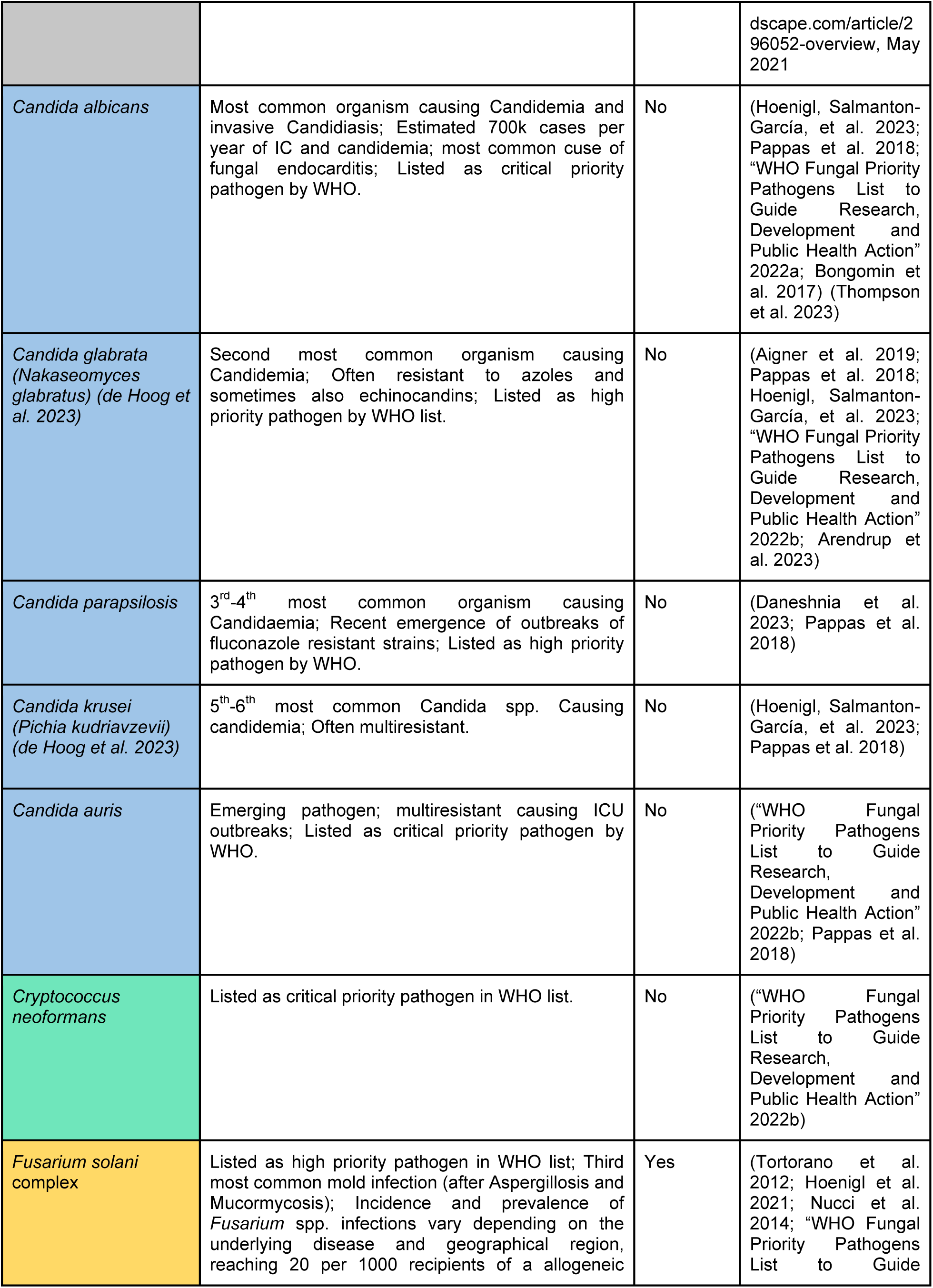
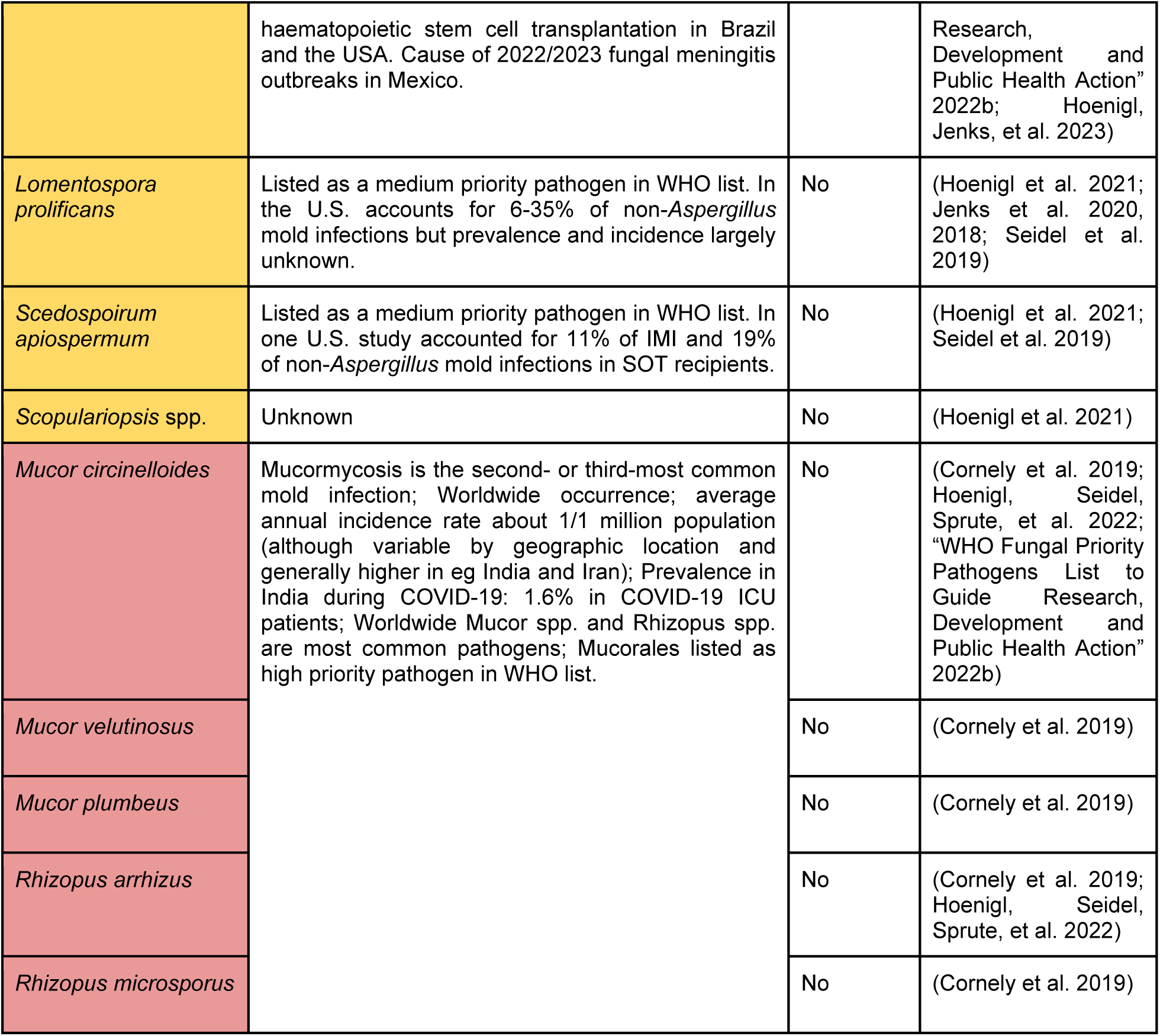
Clinically relevant fungi, including rare molds, used to develop universal assay.

However, *Aspergillus* spp. were not consistently amplified by our ITS primers and the efficiency of this region for detection of *Aspergillus* spp. isolates is not optimal. Furthermore, the ITS1-4 region is not sufficient for discriminating between many individual *Aspergillus* spp. And has been shown to not amplify in certain isolates (Fraczek et al. 2019; “Target Genes, Primer Sets, and Thermocycler Settings for Fungal DNA Amplification” 2022, “Evaluation of Nucleic Acid Sequencing of the D1/D2 Region of the Large Subunit of the 28S rDNA and the Internal Transcribed Spacer Region Using SmartGene IDNS Software for Identification of Filamentous Fungi in a Clinical Laboratory” 2012) (Supplementary Fig. 2). Since *Aspergillus* spp. is one of the most clinically relevant fungal pathogens in the US but also globally, we next selected an *Aspergillus*-specific primer set targeting the 18S rDNA gene, which harbors species-specific sequence differences (Supplemental Figure. 3).

This primer set was multiplexed with the ITS primer set and the assay was tested for its ability to amplify the 21 species in Table 1. Our *Scopulariopsis* spp. isolates were not consistently amplified which has been observed previously (“Evaluation of Nucleic Acid Sequencing of the D1/D2 Region of the Large Subunit of the 28S rDNA and the Internal Transcribed Spacer Region Using SmartGene IDNS Software for Identification of Filamentous Fungi in a Clinical Laboratory” 2012), while *Scedosporium apiospermum* isolates produced variable melts indicating multiple organisms (Supplementary Fig. 4), and neither of these were added to the final database. Now, 19 amplified and sequenced in qPCR and produced reliable melt curve signatures in U-dHRM. Figure 2 shows the digital melt curve signatures for each organism and their average curve in black.

**Figure 2.**
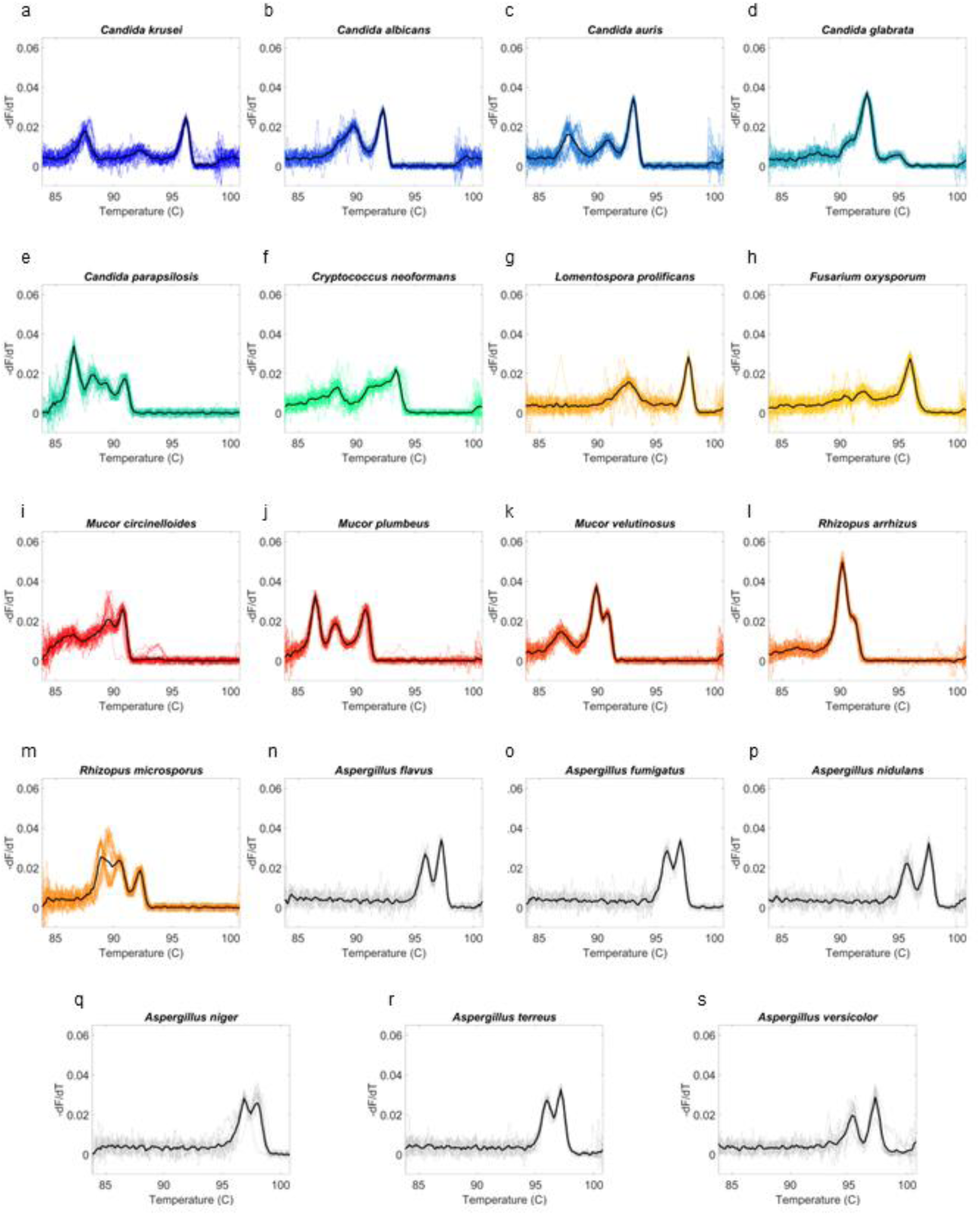
U-dHRM curves for 19 organisms. Asp and ITS primers were multiplexed in U-dHRM and the assay was tested for the detection and melt-based discrimination of 19 organisms: (A-E) *Candida* spp., (F) *Cryptococcus* spp., (G) *Lomentospora* spp., (H) *Fusarium* spp., (I-K) *Mucor* spp., (L-M) *Rhizopus* spp., (N-S) *Aspergillus* spp.. Yeasts are blue/green, molds are orange/red/black.

Next, we conducted analytical validation studies on *Aspergillus* spp. and *Candida* spp. to assess the overall detection capability of the assay in combination with sample preparation starting from a real sample matrix. Mock samples were created by spiking whole organisms into pig BALF over a concentration range of approximately 1×10^5^- 1×10^0^ CFU/mL and no spike controls. Host DNA depletion and pathogen DNA extraction was carried out using MolYsis Complete5 per manufacturer’s instructions. Then, the extracted DNA was loaded onto dPCR chips with the multiplexed Asp+ITS universal fungal assay and amplification was performed prior to dHRM analysis (Supplementary Fig. 5). Fungal melt curve counts showed good linearity of quantification (r2=0.99) for *Candida* and *Aspergillus* spp. (Supplementary Fig. 6A,B). However, *Aspergillus* spp. detection was 10-fold lower than expected and Candida detection was 10-fold higher than expected, based on spore counting and plating. To test if this difference could be attributed to *Aspergillus* spp. being more difficult to lyse or whether it reflected assay sensitivity differences, we conducted *Aspergillus* spp. DNA dilution series experiments. This showed that the assay alone maintained high linearity of detection down to ∼10 copies/chip, or 25 pg/mL (Supplementary Fig. 6C).

### Database generation and algorithm training

To determine whether fungal organism digital melt curves (Fig. 2) could be reliably and automatically recognized by a ML algorithm, a database of >196,000 curves comprising biological and technical replicates n ≧ 3 for each of the 19 pathogens was generated on dPCR chips. Fig. 3 A-B depicts the ML flowchart comparison for testing database curves versus clinical unknown or novel curves. The classification performance of a ML algorithm that combines dynamic time warping and Euclidean distance based metrics was assessed in cross-validation studies (“Weighted kNN and Constrained Elastic Distances for Time-Series Classification” 2020).

**Figure 3.**
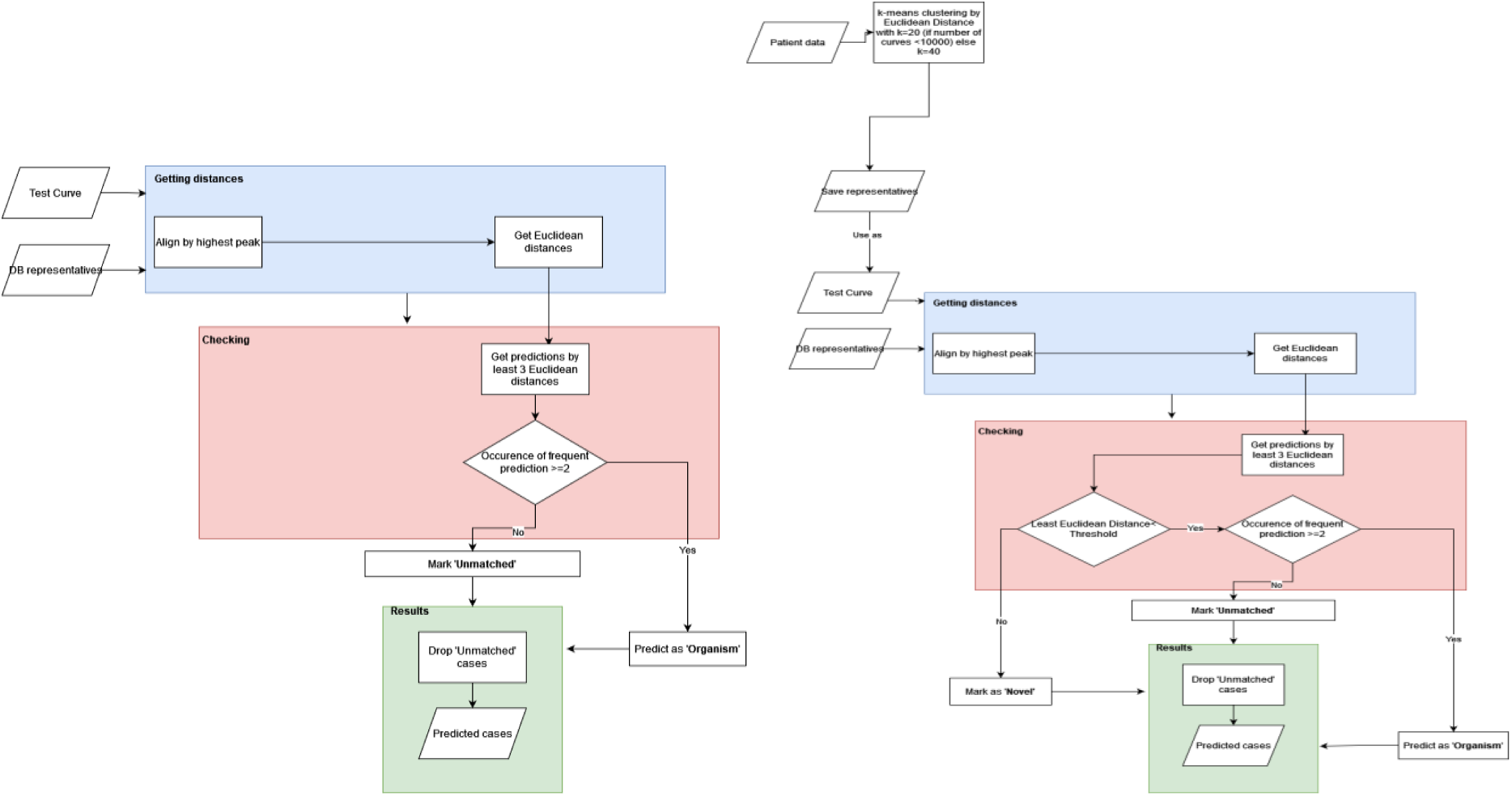
Machine learning process. a) Flowchart for database sample testing b) Flowchart of patient samples testing differentiating between database classification and novelty detection

Recall was assessed and plotted as a confusion matrix in Fig. 4A. This revealed that *Aspergillus* spp. were not reliably discriminated within the genus, while all other species were reliably classified. Among *Aspergillus* spp., cross-validation showed that an overall accuracy (F-score, a combination of precision and recall) of about 60% was achieved (Supplementary Table 1). This can be explained visually by overlaying representative curves from each species, which are quite similar (Fig. 4B), due to few sequence differences (Supplementary Fig 4). An overall accuracy of 86% was achieved across the 19 organisms with *Aspergillus* spp. treated as separate classes (Supplementary Table 2). Grouping *Aspergillus* spp. into a single class (Fig. 4C) at the genus level resulted in a significant improvement in the F-score for *Aspergillus* spp. (90%, Supplementary Table 3), and an overall accuracy for all classes of 97% was achieved. The associated confusion matrix (Fig. 4D) shows only 3.4% misclassification overall (5059/150752), with the most occurring between *M. circinelloides* and *Aspergillus* spp. when *Aspergillus* genus is the true class (7.2%, 768/10657). Representative melt curves for each organism class are shown in Fig. 4E.

**Figure 4.**
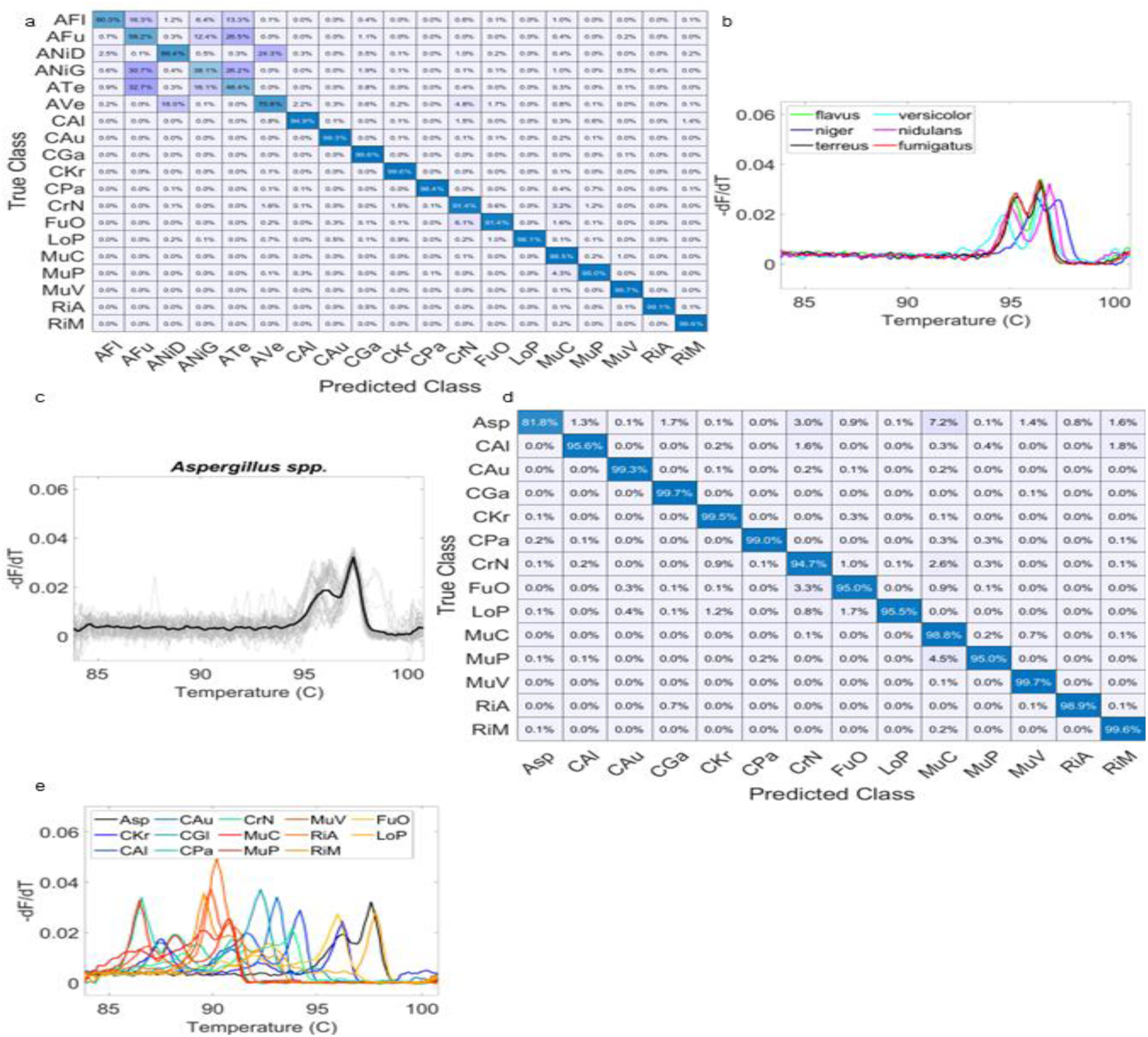
Machine Classification Performance on Fungal Melt Curve Database and Curves. a) confusion matrix with individual *Aspergillus* spp. b) *Aspergillus* spp. average curves overlap c) Grouped *Aspergillus* spp. average curve overlap. d) confusion matrix with grouped *Aspergillus*. e) Average curves of Grouped *Aspergillus* genus and all average curves of 13 other spp.

### Clinical BALF Sample Analysis

#### Overall performance for pathogenic mold detection

U-dHRM achieved an average of 97% fungal organism identification accuracy and a turn-around-time of 4hrs. In total, 73 remnant banked BALF samples that were collected due to suspicion of IMI were analyzed by U-dHRM and compared to clinical diagnostic classifications (Fig. 5). U-dHRM detected pathogenic molds (*Aspergillus, Mucorales, Lomentospora* and/or *Fusarium* spp.*; ≧1 curve* or 11 CFU/mL) in 73% (53/73) of all the samples (Fig. 5A). In addition *Candida* spp. were detected in 88% (64/73) of all samples, while 12% (9/73) had non-*Candida* yeasts as well. We note that there was no apparent association between human β-actin Ct and concentration of fungi detected by U-dHRM or BALF sample volume and concentration of fungi detected by U-dHRM (Supplementary Fig. 7). In 19% (14/73) of samples, mixtures of pathogenic molds were detected (Fig. 5A). Examples of curve signatures detected by U-dHRM and identified by ML in the clinical BALF samples and their closest matching database curve are shown in Supplementary Fig. 8. Of the samples considered positive for IMI, U-dHRM detected pathogenic molds in 73% (1/1 proven; 17/25 probable; 4/4 putative). In samples that were not classifiable for IMI, U-dHRM detected pathogenic molds in 90% (9/10). However, in samples considered negative or without mycological evidence for IMI, U-dHRM detected pathogenic molds in 67% (1/3 possible; 21/30 no). These samples were considered negative for IMI predominantly because of GM and culture negativity as well as absence of host factors, but nonetheless, they were collected due to some clinical suspicion of IMI. These results suggest that U-dHRM has good sensitivity for IMI, as defined by current diagnostic criteria, when host risk factors are also considered. Specificity was optimized by requiring the number of pathogenic mold curves detected in a sample to be *>8* and sample volume to be 1mL, which resulted in a subset of 43% detection in criteria-matching positives (6/14), 50% (5/10) in not classifiable, and 0% detection in negatives (0/21).

**Figure 5.**
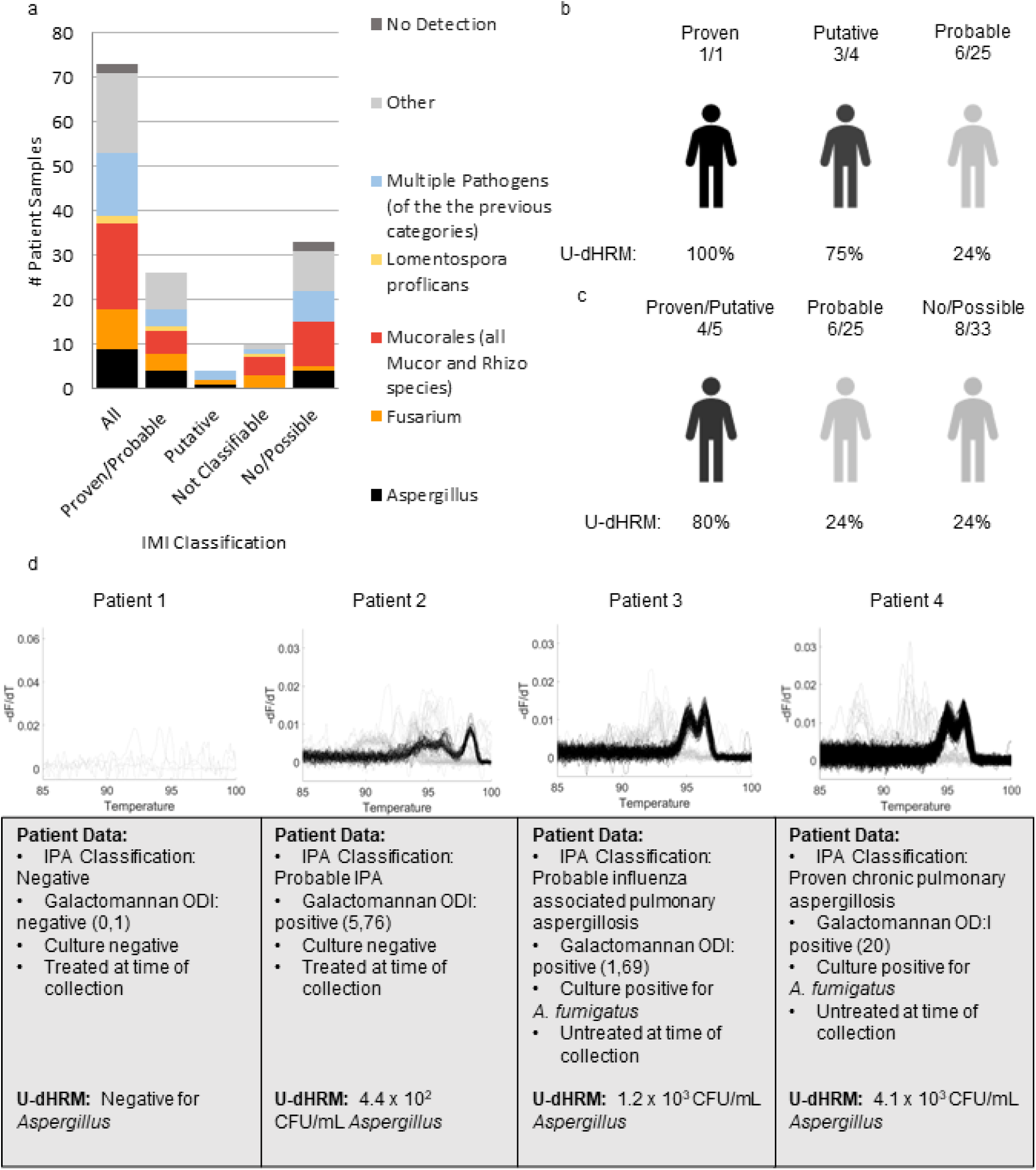
U-dHRM Pathogen Detection Statistics in Patient Samples. a) Pathogen distribution by IMI diagnosis classification. Others are defined as yeasts in the U-dHRM database or unknown novel organisms b) U-dHRM detection of *Aspergillus* in suspected IMI cases ordered left to right by decreasing confidence of suspicion by IPA classification. c) U-dHRM detection of *Aspergillus* in combined highest, medium, and low suspicion. d) Examples of Aspergillus Detection by U-dHRM in BALF. Concordant *Aspergillus* detection confidence and overlapping ML representatives depicts examples of Asp-ITS U-dHRM detection and ML classification in patients with no IPA, probable IPA (treated at the time of collection and culture negative, and untreated at the time of collection and culture positive) and proven IPA. Correlations with routine mycological test results show that more *Aspergillus* curves were detected in the patient with probable IPA who had both, positive BALF GM and positive culture versus the other patient with probable IPA who had only positive BALF GM.

#### *Aspergillus* detection by U-dHRM compared to culture and GM

A summary of *Aspergillus* spp. detection by U-dHRM compared to clinical diagnostic criteria is shown in Fig. 5B-C. Of all the samples that cultured *Aspergillus* spp., U-dHRM detected *Aspergillus* spp. melt curves in 61% of positives (1/1 proven; 4/9 probable; 3/3 putative), 0% of not-classifiable (0/2) cases or no IPA (0/1 no). Considering only samples from proven, probable and putative cases that were culture+, GM+, antifungal treatment-, U-dHRM detected *Aspergillus* spp. melt curves in 78% (7/9). Examples of *Aspergillus* spp. melt curves from patient samples that correlated with routine mycological test results are shown in Fig. 5D. The highest *Aspergillus* spp. load was detected in the patient with proven IPA, and the second highest load was detected in a patient with probable influenza-associated pulmonary aspergillosis (IAPA).

U-dHRM also detected *Aspergillus* spp. in some samples that did not culture *Aspergillus* spp.: 10% (2/19) probable; 12% (1/8) not classifiable; 28% (8/29) no IPA. In samples that did not culture *Aspergillus* spp., other pathogenic molds were often detected alone or in combination with *Aspergillus* spp.: other molds were detected in 71% of probable and putative cases (12/17); 70% (7/10) not classifiable cases; 67% (22/33) of possible and no IPA cases.

#### Differentiation between *Aspergillus* spp. and *Fusarium* spp. by U-dHRM in GM positive samples

Of all the GM+ samples, U-dHRM detected GM-producing organisms *Aspergillus* and/or *Fusarium* spp. in 54% (21/39). Mixtures of *Aspergillus* and *Fusarium* spp. were detected in 8% (3/39). In GM+/*Aspergillus* spp. culture+ samples, *Aspergillus* spp. alone were detected in 36% (5/14), *Fusarium* spp. alone in 7% (1/14), while both were detected in 21% (3/14). In GM+/*Aspergillus* spp. culture-samples, *Aspergillus* spp. were detected in 12% (3/25), *Fusarium* spp. in 36% (9/25), while both were detected in 0% (0/25). These results are depicted in Fig. 6A-B. An example of multiple pathogen detection including *Aspergillus* and *Fusarium* spp. melt curves from a patient sample is shown in Fig. 6C.

**Figure 6:**
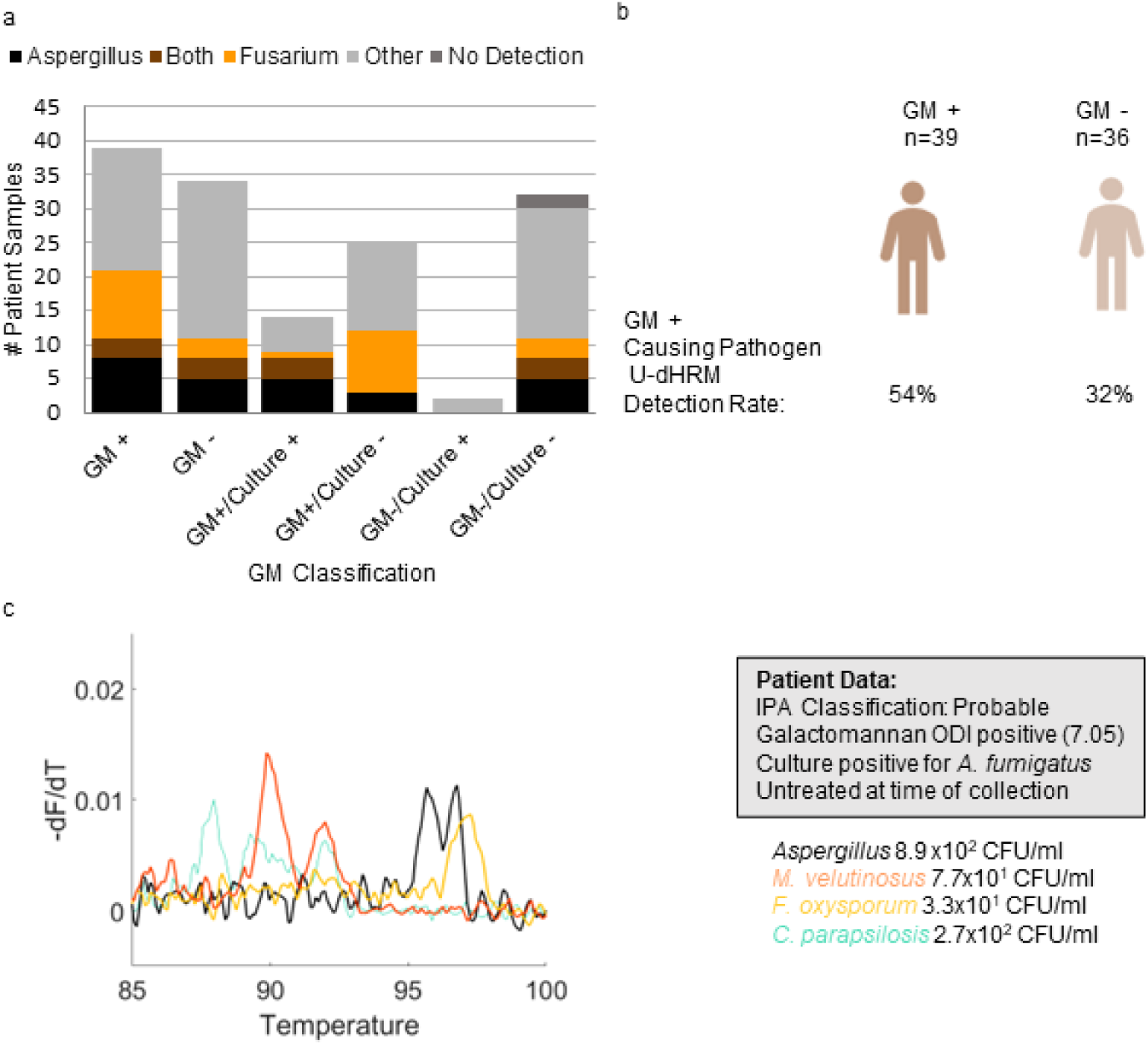
*Aspergillus, Fusarium* multiple pathogen co-detection. a) *Aspergillus* and *Fusarium* detection distribution by GM and culture positivity. Others are defined as yeasts in the U-dHRM database or unknown novel organisms b) U-dHRM detection of GM-producing spp. compared to clinical GM status. c) *Aspergillu*s, *Fusarium*, and *Mucor* codetection example of representative raw clinical curves demonstrating patient BALF pathobiome. Curves are shown for *Aspergillus*, *Mucor*, *Fusarium*, and *Candida* for visualization purposes with the following quantification: 8.9×10^2^ CFU/mL *Aspergillus*, 3.3×10^1^ CFU/mL *F. oxysporum*, and 7.7×10^1^ CFU/mL *M veluntunsosis*, 2.7×10^2^ CFU/mL *C..parapsilosis*, 3.1×10^2^ CFU/mL novel organisms (not shown).

#### Detection of pathogenic molds in the absence of *Aspergillus*

In samples where no *Aspergillus* spp. was detected by U-dHRM, other pathogenic molds were detected in putative 1/1 (100%), probable 11/19 (58%), not classifiable 89% (8/9) cases, possible 33% (1/3), and no IMI 55% (12/22) cases.

#### Mucorales Detection

Fungal pathogens in the *Mucorales* order were detected in 42% (31/73) of all samples. *Mucorales* was detected in 31% (8/26) of the proven/probable IMI, 50% (2/4) putative, 40% (4/10) not classifiable, 33% (1/3) of the possible cases, and 53% (16/30) of the samples classified as no IMI. Under optimal specificity criteria of pathogenic mold curves detected in a sample to be *>8* and sample volume to be 1mL this subset of detection dropped to 15% (2/13) in proven/probable IMI cases, 100% (1/1) of putative cases, 30% (3/10) of not classifiable cases, and no detection in possible cases or samples classified as no IMI.

Co-detection of *≧1* curve for multiple *Mucorales* spp. occurred in 10% (7/73) of samples, with the highest rate in possible 33% (1/3), followed by not classifiable 20% (2/10), no IMI 10% (3/30), probable, probable 4% (1/25), with proven and possible at 0%. Co-detection of *Mucorales* and *Aspergillus* spp. occurred in 11% (8/73) of samples, with the highest rate in putative 50% (2/4) followed by samples classified as no IMI 13% (4/30), proven/probable 8% (2/26), with no co-detection in those classified as possible and those classified as not classifiable. These results are depicted in Fig. 7A-B. An example of co-detection of *Mucorales* spp. including melt curves from a patient sample representing discordant mold diagnosis is shown in Fig. 7C.

**Figure 7.**
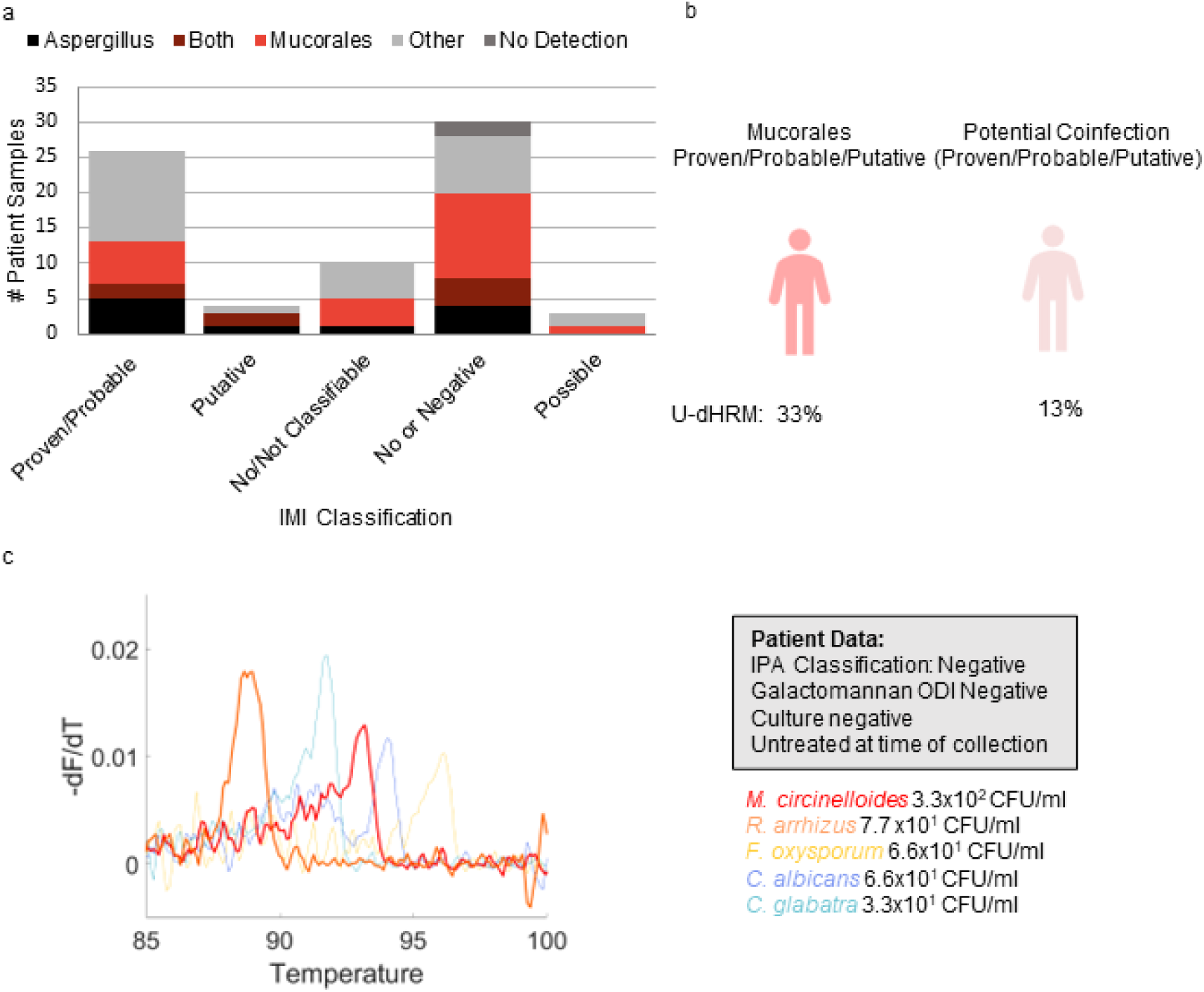
Mucorales detection. a) *Aspergillus* and *Mucorales* detection distribution by IMI diagnosis classification. Others are defined as yeasts in the U-dHRM database or unknown novel organisms b) U-dHRM detection of mucorales and potential co-infection in suspected IMI cases. c) Discordant mold diagnosis example of representative raw clinical curves demonstrating patient BALF pathobiome. Curves are shown for. *Mucor*, *Fusarium*, and *Candida* for visualization purposes with the following quantifications: 6.6×10^1^ CFU/mL *C. albicans* (blue) 3.3×10^1^ CFU/mL *C. glabrata* (blue), 3.3×10^2^ CFU/mL *M. circinelloides* (red), 7.7×10^1^ CFU/mL *R. arrhizus* (orange), 6.6 x10^1^ CFU/mL *F. oxysporum*, and 1.2×10^2^ novel organisms (not shown)

#### Identification of Organisms Generating Novel Fungal Melt Curves

A unique feature of the U-dHRM trained ML algorithm is its ability to automatically detect novel organisms by their distinct melt curve shapes compared to common pathogen curves represented in the database (see Methods). 96% (70/73) of the BALF samples tested produced melt curves that confidently matched to the U-dHRM database of common pathogens. However, a few patient samples generated fungal melt curves that did not match the database and were called novel by the algorithm. To identify the organisms generating these curves, a micromanipulator was used to recover individual digital reactions and sequence their amplicons. Fig. 8 demonstrates the application of this new technique to a patient sample where novel melt curves dominated U-dHRM results (Fig. 8A). Custom software was used to determine the XY position of novel curve generating wells (Fig. 8B), and wells were sampled by using a micromanipulator (Fig. 8C) to position a micropipette into the target well (Fig. 8D) and extract the reaction containing novel amplicons (Fig. 8E). In this sample, *Trichosporon asahii* and *Sacchromyces cerevisae* (Fig. 8A, dark and light gray curves respectively) were identified. Using the recovered *T. asahii* amplicons as template, U-dHRM was conducted to generate database curves for training the ML algorithm to automatically identify this organism in future samples. Supplementary Table 4 describes other patient samples where novel amplicons were recovered and identified, including potentially causative pathogens and commensal yeasts *Pneumonocystis jirovecii*, *Sporobolomyces salminocolor*, *Sacchromyces cerevisae*, *Epicoccum nigrum*, and *Candida inconspicua*. This process avoids the need to culture amplify isolates, which is important considering the low sensitivity of BALF culture and potential fastidiousness of novel organisms. Additionally it will further expand the database while limiting the occurrence of future unidentifiable melt curves, thus minimizing the need for future sequencing, which in turn affects the turnaround time (TAT).

**Figure 8.**
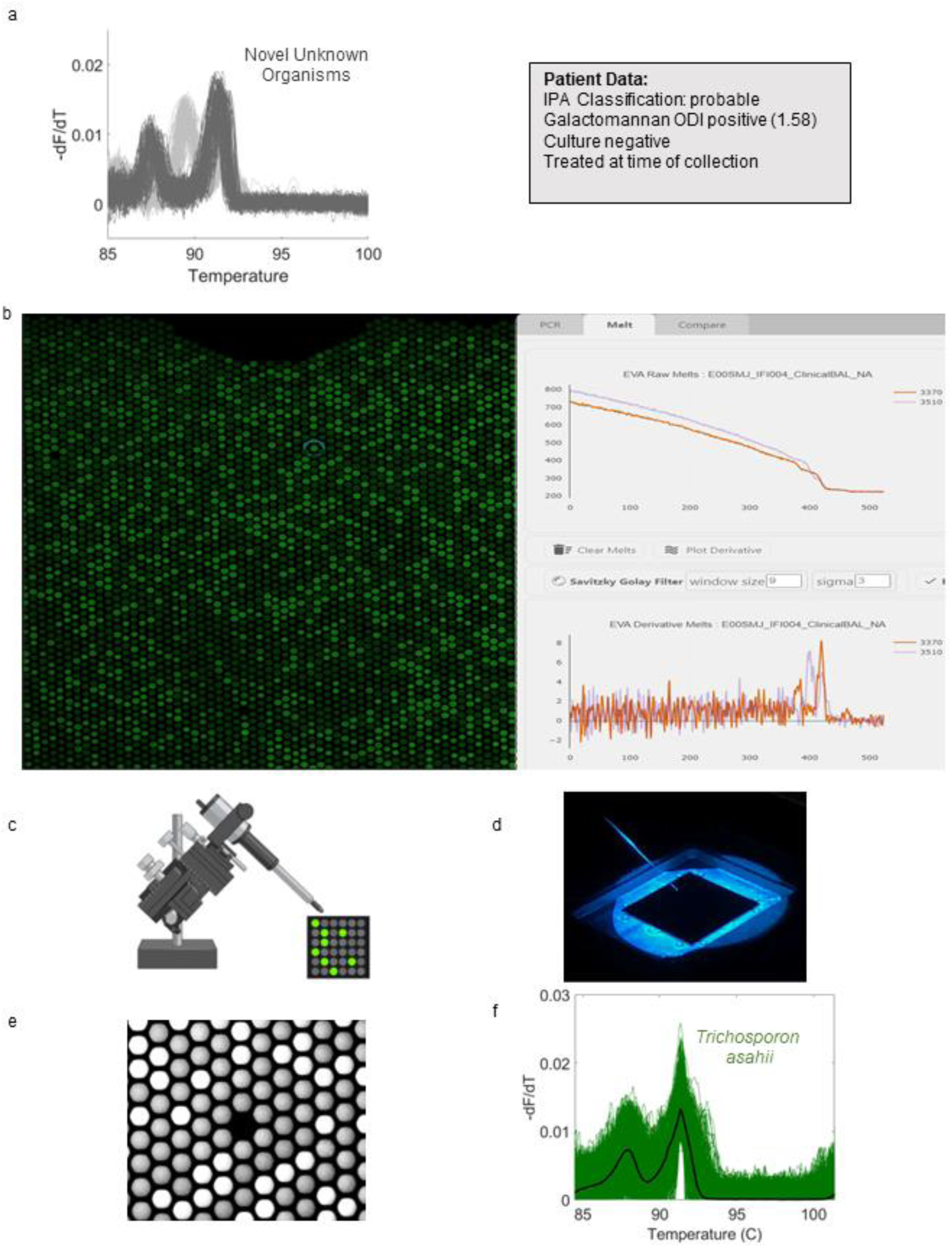
Novel Melt Curve Identification and Algorithm Retraining. a) Novel fungal melt curves *Trichosporon asahii* (dark gray) and *Sacchromyces cerevisae* (light gray) identified by ML for patient sample IFI 004. Diagnostic information for this patient is shown in the adjacent gray box. b) Screenshot of the Melio Melt Inspector software used to find the specific XY location of the wells on-chip harboring novel amplicons. c-d) Schematic and photograph of the micromanipulator positioning a micropipette into the target well for novel amplicon collection. e) Fluorescent micrograph of chip after micropipette extraction of reaction from the target well. f) U-dHRM melt curves generated by re-amplification of the novel amplicon for database expansion and training of the ML algorithm.

## Discussion

Evaluating the performance of new diagnostic tests for IMI is difficult due to the limitations of comparing these new modalities to imperfect gold standard clinical tests and the rarity of autopsy proven IMI, and the ongoing debate over the accuracy of diagnostic classifications (Arvanitis et al. 2014). U-dHRM results were not particularly well correlated with GM positivity or *Aspergillus* spp. culture results - neither of which correlated well with each other - but did demonstrate strong agreement with clinical mycology tests in general and showed good sensitivity for IMI, as defined by current diagnostic criteria, when host factors were also considered. A lack of concordance between GM positivity and U-dHRM detection (or *Aspergillus* spp. culture results) could arise from organism clearance when antigen levels are high or the presence of organisms before antigens are developed during active growth. Importantly, in cases where GM positivity did not correlate with *Aspergillus* spp. detection by culture, U-dHRM results occasionally provided potential explanations by detecting other GM producing organisms such as *Fusarium* and *Trichosporon* spp. (“Serology Anno 2021—fungal Infections: From Invasive to Chronic” 2021).

With BAL culture showing limited sensitivity for detecting pulmonary fusariosis (Hoenigl et al. 2021), *Fusarium* spp. infections resulting in GM positivity can lead to a false diagnosis of probable IPA and incorrect or inadequate antifungal treatment for these highly resistant pathogens. In San Diego *Fusarium* spp. Have been shown to be a frequent cause of rare mold infections (Jenks et al. 2018). U-dHRM had higher detection of *Fusarium* in GM+ samples that did not grow *Aspergillus* spp. in culture. Also, the ability of U-dHRM to detect multiple common pathogens, even in mixtures, has potential to identify mixed infections and improve treatment decisions. For example, in a patient classified as probable for IPA with positive GM and *Aspergillus* spp. Culture results, U-dHRM detected *Aspergillus* spp. in concordance with these results, but also detected *F. oxysporum,* and *M. velutinosus* at similar abundances (8.9×10^2^ CFU/mL *Aspergillus*, 3.3×10^1 *F. oxysporum*, and 7.7×10^1^ CFU/mL *M. velutinosus*, 2.7×10^2^ CFU/mL *C. parapsilosis*). While at the time of BALF collection this patient had not received antifungal treatment, the patient subsequently received treatment for IPA with voriconazole (which likely covered *F. oxysporum* but not *M. velutinosus*) and passed away within a week, with no autopsy performed. In this case, U-dHRM results may have influenced treatment to include antifungals targeting *M. velutinosus*. In another example, U-dHRM detected a mixture of different *Mucorales* spp. in a patient with suspected IMI but negative GM and *Aspergillus* spp. culture results. Of note, one of the species detected, *M. circinelloides*, commonly shows higher MICs against isavuconazole and posaconazole, complicating therapy (Badali et al. 2021). While IPA can be diagnosed with the presence of host factors, clinical symptoms, radiological findings, and mycological evidence of *Aspergillus* either in culture or by detection of GM, other IMIs can mimic the clinical presentation of IPA, with mycological evidence mostly limited to insensitive culture or histology. As a prominent example, mucormycosis diagnostics is particularly challenging (Cornely et al. 2019). Pulmonary mucormycosis remains one of the most common non-*Aspergillus* mold infections in many US centers, and has been globally and particularly in India on the rise as a complication in COVID-19 patients(Hoenigl, Seidel, Carvalho, et al. 2022). There is hope on the horizon with Mucorales PCR now starting to be implemented in some clinical centers (Guegan et al. 2020).

While U-dHRM detected *Aspergillus* spp. in 61% (9/13) of culture positive samples from patients with IPA, the method also detected *Aspergillus* spp in 19% (11/58) of culture negative samples, The presence of viable but non-culturable organisms may explain the this finding. Molysis sample processing upstream of U-dHRM analysis utilizes selective lysis, DNase, and filtration steps to degrade host and cell-free DNA and enrich for intact organisms, which allows U-dHRM to detect organisms that are intact but may not grow in culture.

The ability of U-dHRM to detect novel fungal organisms also demonstrated diagnostic value in this patient cohort. Several patient samples contained more novel melt curves than curves from common pathogens. The ability to recover these amplicons for same-day Sanger sequencing enabled the fast identification of emerging pathogens of clinical significance. In one case, using this method resulted in the identification of *T. asahii* as the dominant organism in the BALF of a patient classified as probable for IPA with positive GM and negative culture who had already received 42 days of micafungin. U-dHRM did not detect *Aspergillus* spp. In that patient. *T. asahii* is resistant to micafungin and can cause positive GM. It is an emerging pathogen that is rarely identified in clinical practice but often causes fatal infections in immunocompromised individuals due to being misdiagnosed as other types of fungal infections and because of its resistance to many front-line antifungals (Li et al. 2020). This particular patient was never diagnosed with or treated for *T. asahii* and passed away, suggesting that U-dHRM could have provided critical diagnostic value with high impact for this patient.

Overall, the performance of U-dHRM suggests that it could represent a promising advance in molecular pathogen detection strategies for IMI. Previously, broad-based qPCR followed by sequencing has shown promise for improving the detection of rare molds, but this approach is recommended *only* when fungal elements are seen by histopathology due to sensitivity limitations (Donnelly et al. 2019). Also, the presence of multiple fungal species can lead to the detection of only the dominant species or failed detection altogether(Kidd et al. 2019; Zeller et al. 2017). U-dHRM distinguishes itself by implementing broad-based PCR in a higher sensitivity dPCR format. Implementation of melt analysis in a digital format enables identification and counting at the single genome level, even in polymicrobial samples, and eliminates template amplification competition and efficiency biases. Also, this format allows extensive melt curve training data to be rapidly generated, unlocking the power of machine learning through big data for automated melt curve identification to rapidly identify and quantify the sequences of all the common pathogens in the sample individually. Only novel organism curves of high abundance warrant interrogation by sequencing, saving time and expense. U-dHRM technology allows for a broader snapshot of the patient pathobiome, including more sensitively detecting and discriminating causative species. The quantitative nature of U-dHRM results also highlight the potential for monitoring over time to track mixed infections, measure effectiveness of therapies, and aid in discriminating between true infection (growth) versus colonization (stasis).

## Limitations and Future Work

Based on total curve counts per chip and Poisson theory, we estimate that approximately 10% of samples (7/73) had 1.6% of total wells with double occupancy. So multiple organism curves could have overlapped in these wells, which may generate multiplexed curves that would be called novel. Running U-dHRM on a dilution of these samples overcomes this challenge. ML could also be potentially trained on combined melt curves in multiple occupancy wells. Also, *Aspergillus* spp. curves were not reliably differentiable, indicating that the sequence diversity of the *Aspergillus* specific amplicon generated by the selected primers was not sufficient. Future studies should re-engineer the assay to ensure sufficient sequence diversity to yield distinguishable melt curve shapes. A curve number cutoff may need to be implemented for better specificity, potentially establishing different cutoffs for specific pathogens or commensal organisms. The sample processing may have also contributed to some discrepancies, since culture was conducted at the time of sampling but U-dHRM was conducted after samples had been frozen and stored for up to 8 years. Freezing may have led to organism lysis prior to sample preparation, which could contribute to missed detections by U-dHRM, explaining some of the negative results in patients with prior *Aspergillus* detection by culture. Analytical study results also suggested that the lysis step prior to U-dHRM could also be improved to facilitate higher sensitivity for difficult to lyse organisms like *Aspergillus* spp.(Scharf et al. 2020).

In conclusion, the promising performance and speed of U-dHRM and its ability to simultaneously identify and quantify clinically relevant mold pathogens in polymicrobial samples as well as detect emerging opportunistic pathogens may provide information that could aid in treatment decisions and improve patient outcomes. Future studies would ideally be run on freshly obtained BALF samples instead of remnant banked samples, and would also have BALF PCR results as well as concurrent blood samples in order to provide a molecular comparator to help overcome the limitation when comparing against GM and culture, and to help discriminate angioinvasive infections (Mah et al. 2023). Sampling from timepoints before and after IMI classification would aid in evaluating U-dHRM’s diagnostic power compared to the gold standard tests, diagnostic classifications, and response to treatment. While our study has shown the potential of this method to aid IMI diagnosis, all these measures could also help to establish a reliable cutoff for improving specificity for infection versus colonization and thereby accuracy.

## Acknowledgements

We acknowledge and thank MelioLabs for the use of their MeltRead Platform and for their assistance in data analysis with their ML algorithm. We would like to thank Dr. Sarah Reed for providing fungal isolates. We would like to thank Dr. Pedro Cabrales from UCSD for use of his micromanipulator and Julian M. Jimenez from the Fraley lab for work on adapting the micromanipulator for reaction recovery.

## Ethics Statement

The study protocol and all study-related procedures were approved by the Human Research Protections Program at UCSD.

